# Profiling of mature stage human breastmilk cells identifies host-defense lactocyte sub-populations

**DOI:** 10.1101/2021.10.04.463125

**Authors:** John P. Gleeson, Namit Chaudhary, Rose Doerfler, Katherine C. Fein, Trish Hredzak-Showalter, Kathryn A. Whitehead

## Abstract

Breastmilk is chock-full of nutrients, immunological factors, and cells that aid infant development. Maternal cells are the least studied breastmilk component, and their unique properties are difficult to identify using traditional techniques. Here, we characterized the cells in mature stage breastmilk from healthy donors at the protein, gene, and transcriptome levels. Holistic analysis of flow cytometry, qPCR, and single cell RNA sequencing data identified the predominant cell population as epithelial with smaller populations of macrophages and T cells. Two percent of epithelial cells expressed four stem cell markers: SOX2, TRA-1-60, NANOG, and SSEA4. Furthermore, milk contained six distinct epithelial lactocyte sub-populations, including three previously unidentified sub-populations programmed towards host-defense and intestinal development. Pseudotime analysis delineated the differentiation pathways of epithelial progenitors. Together, these data define healthy human maternal breastmilk cells and provide a basis for their application in maternal and infant medicine.

## Introduction

Although human infants are born immature (*1, 2*), their diet of breastmilk enables continued organ growth and development postpartum, particularly within the intestine (*3*). Human milk contains growth factors, immunomodulatory components, and bacterial and maternal cells. A substantial body of research has delineated the role of many nutritional components of milk, including proteins, fats, and sugars, with a primary goal of creating formula that recapitulates breastmilk (*4*). There is also considerable interest in bacterial cells present in the maternal milk microbiome, which help to establish the infant gut microflora (*5*). By contrast, despite their abundance (~10^6^ cells per ml), the biology of breastmilk cells and their potential role in infant development remains poorly understood.

The first step in determining the physiological relevance of breastmilk cells is to identify and characterize them. Various breastmilk cell types have been described to some degree in the literature, including lactocytes, immune cells, myoepithelial cells, and stem cells (*6*). Immune cells are the dominant population in early-stage milk (colostrum; <1 week after birth). Whereas, in mature stage milk from healthy mothers (>2 weeks postpartum), the majority of cells are lactocytes, which are terminally differentiated milk-producing epithelial cells (*7*). Multipotent stem cells have also been identified in breastmilk based on the expression of several stem cell gene markers, including p63 and KLF4 (*8, 9*). These cells were also differentiated into all three germ layers (*10*), a unique characteristic of pluripotent stem cells (*11*).

Although the roles of the maternal cells in breastmilk are not yet clear, there is early evidence that they may participate in several vital functions. For example, bulk RNA sequencing studies showed that the maternal cells in colostrum express a high level of non-nutritive genes (iron-binding, triglyceride catabolism, and protein digestion). Dominant gene expression then shifts in mature stage milk to the two genes responsible for producing milk proteins – α-lactalbumin and β-casein – in mature stage milk (*12*). Establishing baseline concentrations and gene expression profiles for distinct breastmilk cell types might also aid the development of diagnostics for the detection of cancer in the breastfeeding person (*13*).

Additionally, there is a report that maternal stem and immune cells exit the GI tract of the infant and functionally integrate into the infant’s organs, similarly to immune cells (*14*). These data open the possibility that maternal cells play a role in organ development and passive immunity. For example, necrotizing enterocolitis, an intestinal disease associated with the use of formula, affects ~7% of premature infants and leads to 1,000-2,000 infant deaths a year in the US (*15, 16*).

Given this mounting evidence that maternal breastmilk cells are involved in infant development and may have properties amenable to cellular therapies and diagnostics (*17*), it is imperative that milk cells are clearly and comprehensively characterized. To that end, we were motivated to conduct a three-pronged analysis of milk cells with the broadest applicability to nursing people worldwide – fresh, mature-stage milk cells from healthy donors. The experiments reported here were designed to address the limitations of previous milk cell studies.

Specifically, numerous studies relied exclusively on flow cytometry to identify distinct milk cell populations (*10*). This is problematic because of variable fluorescent antibody quality and unavailability of antibodies for novel cell types in milk. Another limitation is that maternal breastmilk cells are sometimes frozen after collection and before analysis (*13*). This can be problematic because some populations of breastmilk cells do not survive the freezing process; thus, analysis biases against cell populations that are not sufficiently robust.

Through this work, we have circumvented these challenges by analyzing fresh maternal breastmilk cells by flow cytometry, quantitative real time PCR (qRT-PCR), and single cell RNA sequencing (scRNA-seq) simultaneously. Although there is one other study that characterized breastmilk cells using scRNA-seq, it was performed on frozen breastmilk samples from women with gestational diabetes (*13*). To our knowledge, our study is the first of its kind, as it holistically describes the complete population of maternal breastmilk cells from healthy women. Because most lactating mothers are classified as healthy, these findings provide a broadly applicable reference dataset for maternal breastmilk cells and lay the foundation for future studies on the role of these cells in infant development and their potential as therapeutic interventions.

## Results

### Epithelial lactocytes are the most abundant cell population in breastmilk

For our initial analysis, we collected freshly expressed breastmilk from donors to characterize the maternal cell populations using flow cytometry and RT-qPCR (Fig. 1). Although recruitment was open to all donors regardless of postpartum age, we received donations only of mature stage milk (≥ 4 weeks, averaging 32 weeks post-partum; Table S1) (*12*). Following the cell isolation process as previously reported (*10*), we assessed viability by flow cytometry using a fluorescent live-dead stain and observed cell viability of 79.2% ± 6.9% (Fig. S1; *N* = 10). The traditional Trypan Blue cell viability assay reported a high degree of false positives due to the presence of milk fat globules; this assay should be avoided with breastmilk cells.

**Fig. 1.**
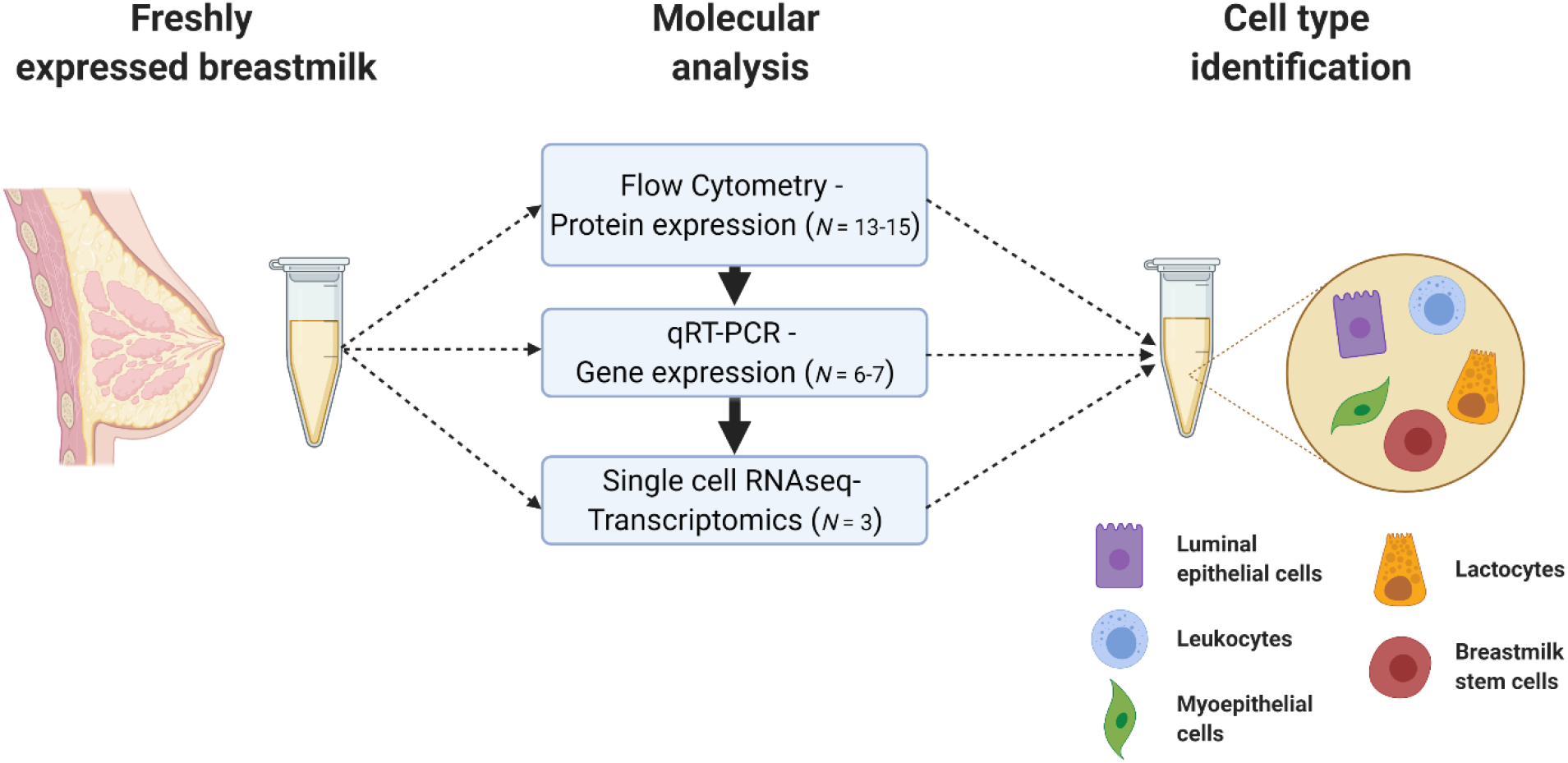
Maternal cells in breastmilk were identified by protein, gene, and transcriptome analyses. Mature-stage breastmilk from healthy donors was assessed for expression of cell marker proteins (e.g. EpCAM, CD45) in freshly isolated samples by flow cytometry. Then, RNA was isolated and used to conduct complementary gene expression analysis by qRT-PCR. Single cell transcriptomics added depth to the analysis and defined the lactocyte population in breastmilk.

To understand the cell populations present, we selected surface markers for flow cytometry analysis based on previous studies: epithelial cells (EpCAM+), immune cells (CD45+), and mesenchymal cells (VIM+) (*6, 10, 12*). We also assayed for lactocytes, which derive from epithelial cells, and thus are EpCAM+ in addition to being CK18+. The selected histograms indicate that the vast majority of breastmilk cells are of epithelial origin (93%), with more modest populations of CK18+, CD45+, and VIM+ cells from healthy donors (*N*=13) (Fig. **2**A). Immune cell numbers increased from ~7% in milk from healthy donor-infant dyads to ~63% and ~36% when the infant (*N*=1) or the mother (*N*=3), respectively, had been sick (Fig. 2B). In the case of illness, the number of donors was too low to run statistical analysis. These flow cytometry data are shown for all donors in (Fig. **2**C) and agree with previous reports, aside from the low percentage of cells expressing CK18 (*10, 13*). However, mRNA expression analysis shows reasonably high expression of *KRT18*, the gene associated with CK18 and lactocytes (Fig. **2**D). Accordingly, we concluded that the lactocyte population observed by flow cytometry was artificially low, likely due to poor binding affinity of the antibody. Gene expression of an immune cell gene, *PTPRC*, and the mesenchymal gene marker, *VIM*, were significantly lower in expression, consistent with the flow cytometry data.

**Fig. 2.**
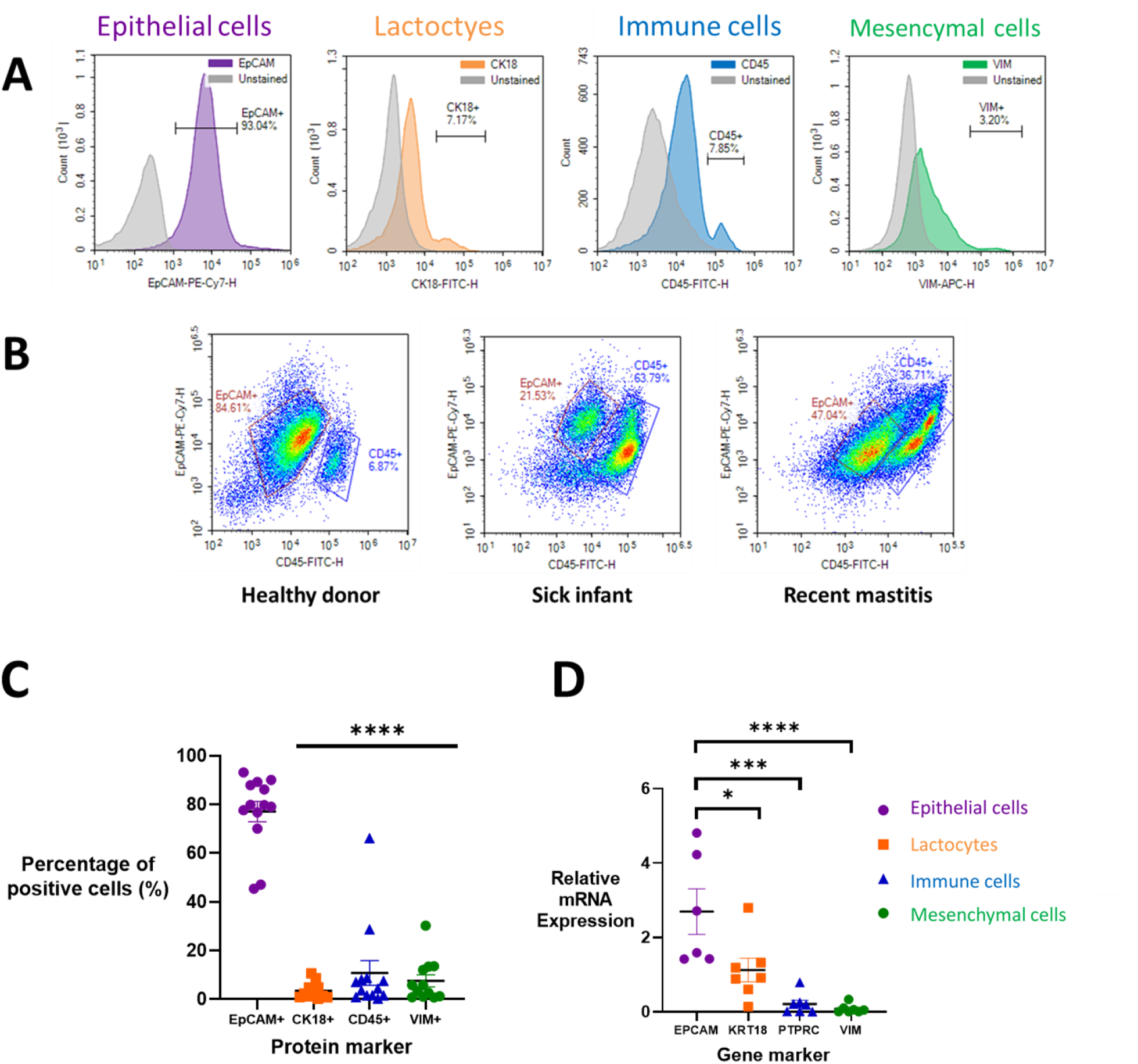
Mature-stage breastmilk from donors contained epithelial lactocytes and a smaller population of immune cells. **(A)** Flow cytometry quantified expression of cell-specific markers: EpCAM+ (epithelial cells), CD45+ (immune cells), CK18+ (lactocytes), and VIM+ (mesenchymal cells). These data suggest that the CK18 antibody bound poorly to lactocytes, and likely underestimates true lactocyte percentages. (B) The relative cellular composition in breastmilk was affected by the health status of the mother and infant. Representative density plots are shown of EpCAM vs CD45 cell populations in milk from healthy donors (*N* = 10), donors with sick infants (*N* = 1), or donors with recent mastitis (*N* = 3). (C) Flow cytometry analysis of protein markers (*N* = 13) and (D) mRNA expression of gene markers corresponding to proteins in (C) indicate the presence of epithelial lactocytes (*N* = 6-7). Data are shown as mean ± SEM. One-way ANOVA with Tukey’s multiple comparison test; * *P <* 0.05, ** *P <* 0.01, *** *P <* 0.001, **** *P <* 0.0001.

### Stem-like transcription factors are highly expressed in 2% of breastmilk cells

There has been significant interest in breastmilk as a novel source of stem cells. Stem cells can be identified by the transcription factors SOX2, NANOG, and OCT4, which are considered the master regulators in stem cells, and by REX1, SSEA4, TRA-1-60, and KLF4, which are co-regulators of pluripotency (*18*). As such, we asked whether the unknown fraction of epithelial cells in breastmilk were stem-like using flow cytometry and qRT-PCR.

For flow cytometry, we gated on EpCAM+ cells and then measured the percentage of cells expressing SOX2, TRA-1-60, NANOG, and SSEA4 (Fig. **3**A). Although we wanted to assess OCT4 as well, we were limited by the constraints of our flow panel. Ultimately, we deemed it more informative to have two master regulator markers and two pluripotency markers in our analysis. These experiments indicated that most epithelial cells were SOX2+ and TRA-1-60+, but significantly fewer were NANOG+ and SSEA1+ (Fig. **3**B).

We also conducted gene expression analysis by RT-qPCR for the genes encoding SOX2, TRA-1-60, NANOG, and SSEA4: *SOX2*, *PODXL*, *NANOG*, and *FUT4*, respectively. We quantified expression levels relative to the housekeeping gene, *EEF1A*, which was highly and uniformly expressed in all breastmilk cells. Epithelial cells expressed 1,000-fold greater *FUT4* mRNA than the other assessed genes, with decreasing expression of *PODXL* > *NANOG > SOX2* (Fig. 3C). The higher *FUT4* expression is likely due to its role in human milk oligosaccharide synthesis, so expression here may not be solely pluripotency related (*19*). When considered together, these data indicate that ~2% of the epithelial cells were positive for all four transcription factors assessed.

**Fig. 3.**
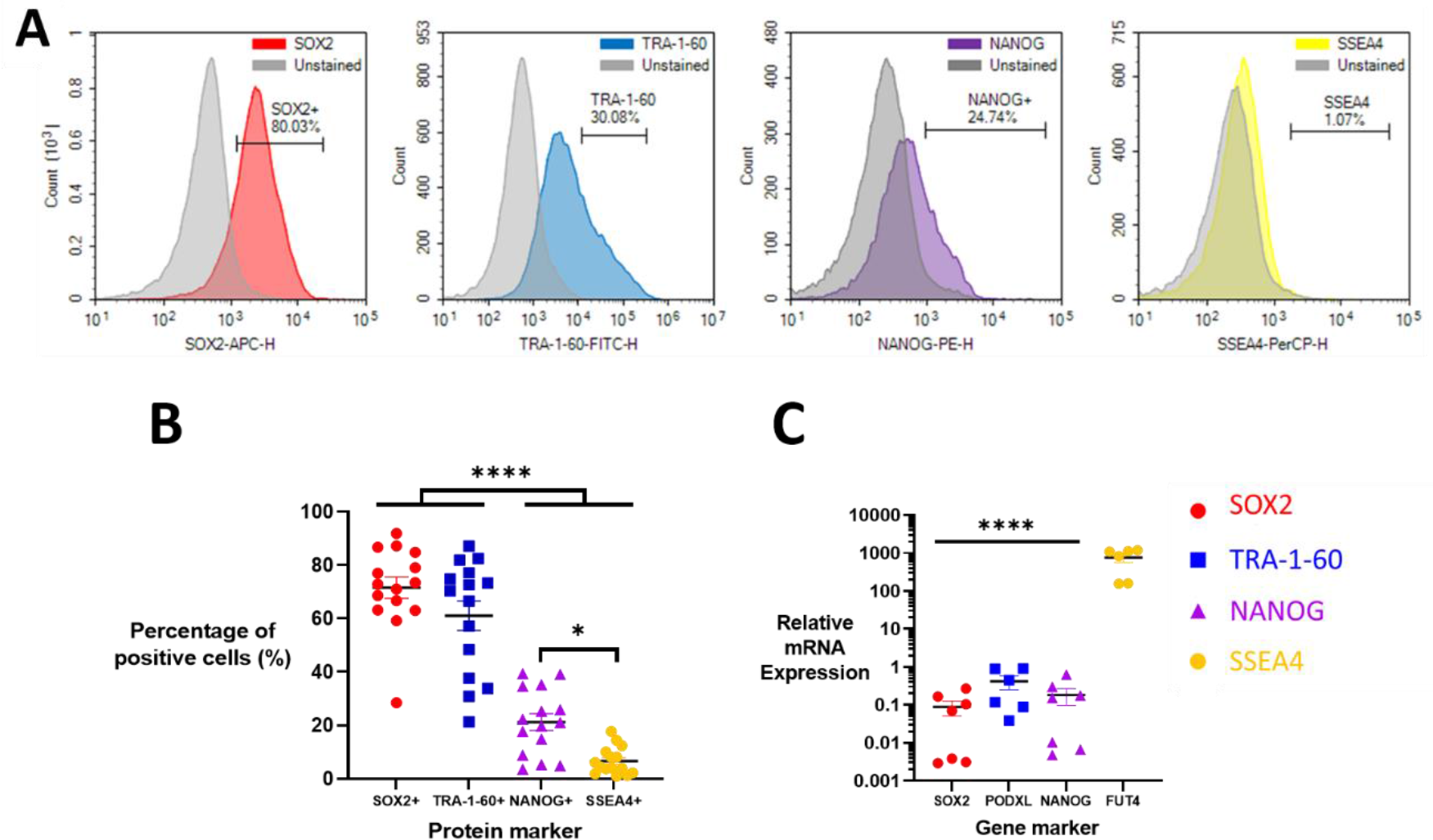
Stem-like transcription factors were expressed in 2% of breastmilk cells. Four stem-like transcription factors were assessed in breastmilk epithelial cells. (A) From flow cytometry analysis, representative histograms of SOX2+, TRA-1-60+, NANOG+, and SSEA4+ stained and unstained samples are shown. (B) Flow cytometry analysis of EpCAM+ cells (*N* = 15) indicated high expression of SOX2 and TRA-1-60 with moderate to low expression of NANOG and SSEA4, respectively. (C) mRNA expression relative to a housekeeping gene was quantified for the genes corresponding to the proteins in (B) (*N* = 7). Data are means ± SEM. One-way ANOVA with Tukey’s multiple comparison test; * P < 0.05, **** P < 0.0001.

### Single-cell transcriptomic analysis reveals unique lactocyte sub-types

Our flow cytometry and mRNA expression data suggest that breastmilk cells are primarily epithelial with a moderate number of lactocytes and a small number of stem-like cells. However, only limited conclusions can be drawn from these two techniques, as the quality of flow data may have been influenced by antibody quality, and bulk gene expression data cannot delineate gene expression patterns in varied cell populations.

To overcome the latter shortcoming, we performed single-cell RNA sequencing to robustly quantify and classify the maternal cell types in milk. For these experiments, we identified six donors with similar cellular profiles by flow cytometry as determined by a first donation. These donors provided samples a second time, and we selected three of those donors (mean 28 weeks postpartum) to advance to scRNA-seq experiments based on sample consistency and cell yield as determined by flow cytometry (Table S1). Upon their third donation, we submitted samples from these three donors within 1-2 hours of expression for scRNA-seq analysis. In total, the transcriptomes of 26,922 maternal breastmilk cells were sequenced, with a median of 1,057 genes per cell.

Gene expression across all three donor samples was dominated by two genes, *LALBA* and *CSN2*, that accounted for > 40% of gene counts (Fig. **4**A). These two genes encode the proteins alpha-lactalbumin and beta-casein, respectively, which are two most abundant proteins in human milk (*20*). The genes identified by scRNAseq were confirmed in a larger donor population by qRT-PCR (*N* = 7), high expression of *CSN2* and *LALBA* with decreasing expression of *CSN1S1 > FTH1 > LYZ*, genes encoding the proteins alpha-S1-casein, ferritin heavy chain 1, and lysozyme, respectively (Fig. **4**B).

To identify the main cell types within the breastmilk cell population that are generalizable across individuals, we performed a combined analysis of the scRNA-seq data using the Seurat and Monocle pipeline (*21*). This analysis identified one large group of cells comprising the contiguous clusters 1 – 7 as well as two smaller independent clusters (8 and 9) based on Uniform Manifold Approximation and Projection (UMAP) clustering (*22*) (Fig. **4**C). Comparison across donors showed that their cluster mappings were reasonably homogenous (Fig. **4**D).

We then examined the differential expression of genes within each cluster (Fig. **4**E), which identified two immune cell clusters, cluster 8 as macrophages (*CD68*+, *ITGAX*+) and cluster 9 as T-cells (*PTPRC*+, *CD3E*+). Although the major cluster (1 – 7) was epithelial (*LALBA*+, *EPCAM*+), there were unique sub-populations that we believe to be previously unidentified in human breastmilk: *FGFBP1*+ lactocytes for intestinal development (cluster 2), fatty-acid synthesizing and insulin-resistant *PTPRF*+ lactocytes (cluster 6), and secretory/chemotactic epithelial cells (cluster 7). Cluster 5 contained the cells that had undergone the least differentiation. These included a small population of progenitor-like cells, defined by high expression of CD55 and CLDN4 (*23*), and a larger population of differentiating cells, which are the precursors of epithelial cells and lactocytes. Finally, cluster 4 comprised apoptotic epithelial cells, and it is unclear what cluster these cells belonged to prior to degradation.

**Fig. 4.**
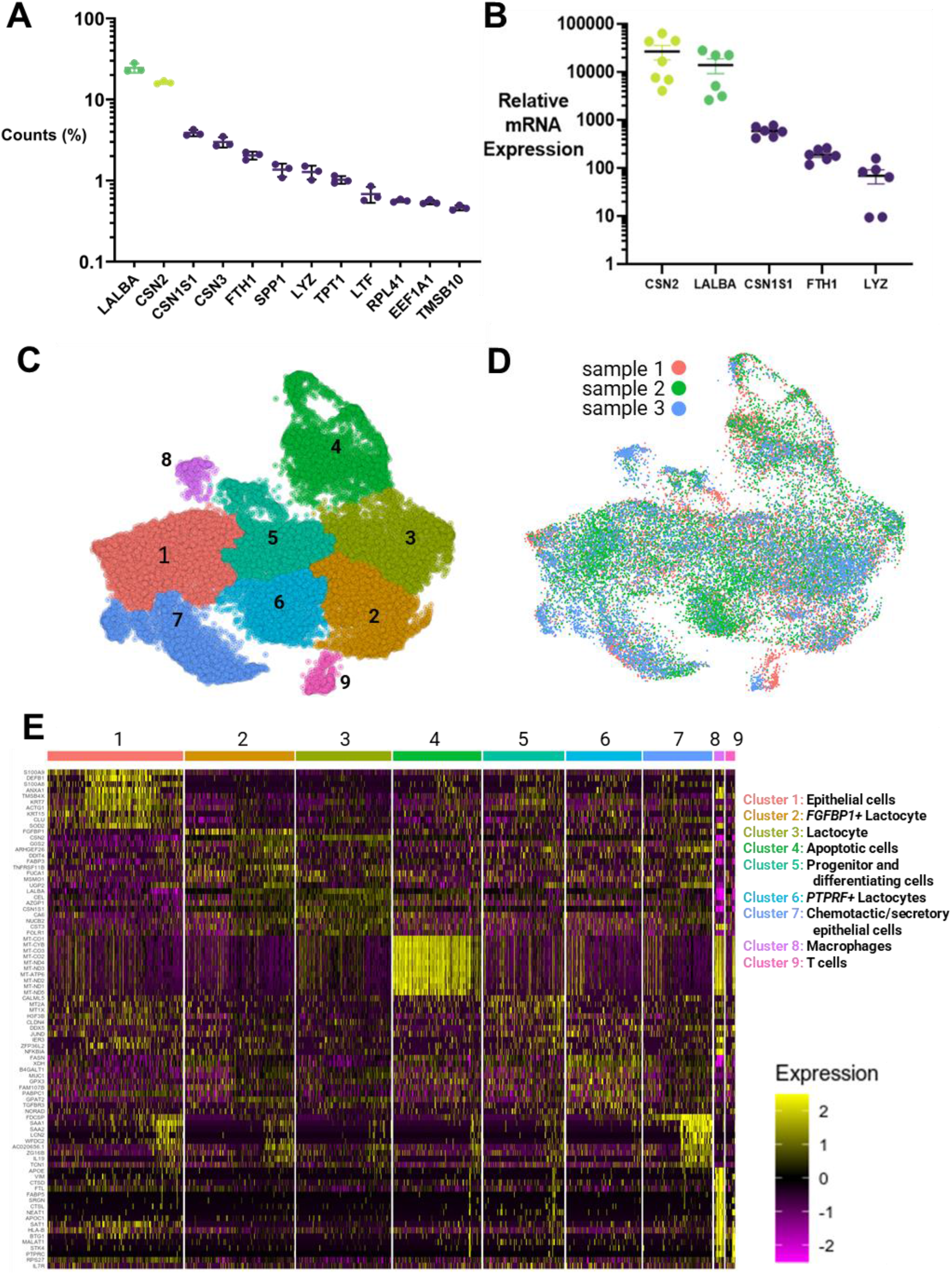
Single-cell transcriptome analysis identified breastmilk cells as primarily epithelial-derived lactocytes, including three novel sub-populations involved in host-defense. (A) Fifty percent of the transcripts produced by mature breastmilk cells from healthy donors corresponded to the milk-producing genes for lactalbumin (*LALBA*) and three types of caseins (*CSN2*, *CSN1S1*, and *CSN3*), N = 3. (B) RT-qPCR quantified mRNA expression relative to *EEF1A* expression for several top genes (*N* = 6). (C) Single cell RNAseq data generated a UMAP plot that identified one major population of cells (Clusters 1 – 7) and two minor populations (Clusters 8 and 9). (D) The UMAP cluster map is consistent across all donors (*N* = 3). (E) Each cluster had unique expression patterns of the top ten marker genes. These patterns were used to identify specific cell types, including three novel types of epithelial lactocytes: *FGFBP1*+ lactocytes*, PTPRF*+ lactocytes, and chemotactic epithelial cells.

To determine the pathway of differentiation of progenitor-like epithelial cells, we performed pseudotime trajectory analysis (Fig. **5**). Fig. 5A shows the differentiation pathways of the breastmilk cells, originating from the most progenitor-like cells, which are part of cluster 5. The starting node is found in the top, deep purple portion of cluster 5. As pseudotime progresses, the cells differentiate along several distinct pathways, with partially and terminally differentiated cells appearing in reddish-orange and yellow, respectively. Fig. 5B shows the spatiotemporal gene expression profiles of nine selected cell markers. High expression of genes associated with progenitor cells or stemness (e.g. CD44, SOD2) occurred earliest in pseudotime and corresponded with cells in cluster 5 (*24–26*), with SOD2 peaking in cluster 1. These progenitor cells briefly expressed *CALML5*, which was exclusively expressed by differentiating epithelial cells, before they diverged into several cell types. Differentiated cells, which appeared in late pseudotime, included mature lactocytes in clusters 2 and 3, lipid and fatty-acid synthesizing cells in cluster 6, and chemotactic/secretory epithelial cells in clusters 1 and 7 (Fig. 5C).

**Fig. 5.**
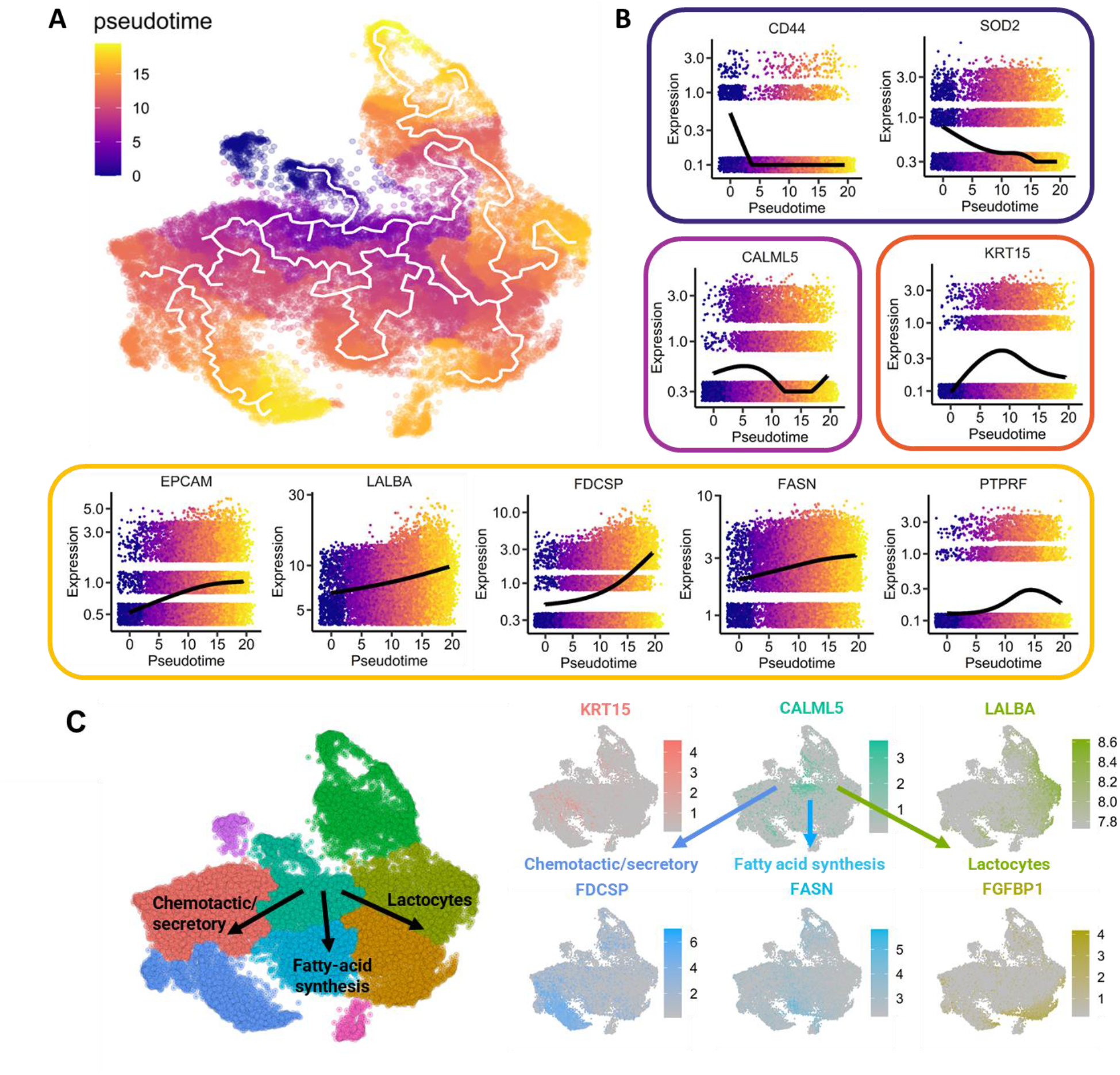
Pseudotime trajectory analysis identified the pathways by which breastmilk epithelial cells differentiate into lactocytes. (A) Pseudotime trajectory analysis determined the pathway of differentiation of progenitor-like epithelial cells (starting node is located in the dark purple region). (B) As pseudotime progressed, the expression of progenitor genes subsided as cells differentiated into mature phenotypes. (C) Based on the pseudotime trajectories and gene expression kinetics, milk cells differentiated along three pathways into chemotactic/secretory epithelial cells, fatty-acid synthesizing cells, and mature lactocytes. Each pathway was associated with overexpression of key genes compared to the other cell types, including *KRT15* (cluster 1), *FDCSP* (cluster 7), *FASN* (cluster 6), *LALBA* (cluster 2), and *FGFBP1* (cluster 3).

Finally, we examined the expression homogeneity of selected markers across seven donors (Fig. S2), two of which were included in our scRNAseq (Table S1). *SOD2* and *CALML5* mRNA were moderately and homogenously expressed (Fig. S2A/B), suggesting progenitor and epithelial cell populations are consistent across donors. By contrast, the chemotactic epithelial marker *FDCSP* had 30-fold and 100-fold higher expression in two donors (Fig. S2D). These data suggest that lactocytes are homogenous across donors, while the specialized sub-types may be enriched for a specific need of the feeding infant, such as intestinal stem cell development (*FGFBP1*+ lactocytes; cluster 2), insulin resistance (*PTPRF*+ lactocytes; cluster 6), or intestinal immune defense (chemotactic epithelial cells; cluster 7).

## Discussion

Through this work, we have provided a detailed characterization of the cellular population in healthy mature-stage breastmilk. We confirmed that most cells were epithelial lactocytes and that ~2% of breastmilk cells are stem-like. Single-cell transcriptomic analysis identified novel sub-populations of lactocytes programmed towards host defense and intestinal development.

Most research on human breastmilk cells has focused on the characterization of stem-like cells present in the milk (*14, 17, 27, 28*). The most thorough of these studies isolated stem-like cells expressing multiple stem-transcription factors (OCT4+, SOX2+, and NANOG+) and differentiated them into all three germ layers (*10*). Unfortunately, this study was unable to determine the percentage of cells positive for more than one marker because the antibodies used in flow cytometry experiments were all FITC-labelled. To our knowledge, we are the first to report that the percentage population of stem-like cells in breastmilk is 2% (SOX2+/TRA-1-60+/NANOG+/SSEA4+) (Fig. 3). Interestingly, single-cell transcriptomic analysis did not identify a cluster of stem-like cells within our breastmilk cell population. This is most likely due to the difference between gene expression and protein translation. However, *SOD2* was expressed well in our samples (Fig. 5B), and this expression has been linked to pluripotency transcription in stem cells (*25*). Further studies are needed to understand the transcriptomic and proteomic profiles of the stem-like cells in breastmilk to determine if they are as useful for regenerative medicine applications, as many authors suggest. Recently, Habowski *et al*. used combined transcriptomics and proteomics to delineate the rapid differentiation of colonic stem cells (*29*). This same type of analysis could potentially determine the stemness and pluripotency of breastmilk-derived cells at a gene and protein level.

Our findings are generally aligned with those reported from the scRNAseq analysis of two donors with gestational diabetes (*13*). Differences observed might be due to the greater number of cells (6x) analyzed in this study, our use of an additional donor, and/or our use of fresh, rather than frozen samples. The use of fresh cells assures that results are not biased towards more robust milk cells that survive freeze/thaw cycles. Further, freeze/thaw processes have been linked to altered gene expression and translation in human primary cells (*30, 31*).

These methodological differences enabled our identification of novel sub-populations: *FGFBP1*+ lactocytes (cluster 2), *PTPRF*+ lactocytes (cluster 6), and chemotactic epithelial cells (cluster 7; Fig. 4E). Regarding cluster 2, *FGFBP1* is a soluble carrier protein that binds to fibroblast growth factor (FGF), which plays a key role in intestinal development and disease regulation (*32, 33*). Cluster 2 also expressed *TNFRSF11B*, which activates the Wnt/β-catenin pathway (essential for intestinal stem cell proliferation) (*34*) and regulates the differentiation of immune sampling M-cells (*35*).

The *PTPRF*+ lactocytes highly expressed *MUC1*, *TGFBR3*, and *PTPRF*. *PTPRF* overexpression in breastmilk cells might be linked to insulin resistant mothers (*12*). *MUC1* secreted in breastmilk binds to the lectin domain of dendritic cells found in infant intestines, preventing pathogenic interactions (*36*). *TGFBR3* plays a key role in activation and inhibition of T and B cells respectively (*37*). In summary, this population may contribute to maternal insulin-resistance signaling and host-defense in the infant.

Finally, chemotactic epithelial cells (cluster 7) had high expression of *FDCSP*, *IL19*, *SAA1*, *WFDC2*, *TCN1*, and *LCN2*. *FDCSP* is considered an ancestral precursor gene to *CSN3*, which produces the milk protein κ-casein (*38, 39*). The other genes with high expression in this cluster appear to confer host-defense, chemokine signaling, and intestinal immune maturation. For example, IL-19 expression was increased in the milk cells of lactating women with mastitis that had been treated with oral probiotics (*40*). The bioactive protein product of *SAA1* in human milk confers protection to the neonate in part by intestinal host-defense and Th17 activation (*41*). Additionally, *WFDC2* contributes to innate immune defenses of the lung, nasal, and oral cavities (*42, 43*), and *LCN2* expression occurs in response to small bowel villous disruption (*44*). Bach and colleagues suggested that mammary epithelial cells should be conceptualized as being part of a continuous spectrum of differentiation, and our results support this (*45*). Overall, our data suggest that the unique lactocyte sub-types identified here by scRNAseq are temporally and uniquely suited to the health and development of mother-infant dyads and strengthen the infants’ immune system during early development.

These findings have provided a genetic fingerprint for the cells in healthy, mature-stage breastmilk and a framework for examining the functionality of these cell types in biologic, prophylactic, and therapeutic contexts. Future work will determine the role of the sub-populations identified here in healthy and sick infants and apply those learnings to the development of novel, oral therapeutic interventions for gastrointestinal and other infantile diseases.

## Materials and Methods

### Breastmilk sample collection and cell isolation

This study was approved by Carnegie Mellon University Institutional Review Board (IRB) under the protocol number – STUDY2019_00000084. Participants were recruited via flyers and social media advertisements in mother and infant groups in the greater Pittsburgh area. All participants provided informed written consent. Pump-expressed mature breastmilk was obtained from each participant at lactation suites at Carnegie Mellon University, placed on ice, and transported to the laboratory immediately after expression for analysis. Breastmilk samples were processed as previously reported (*10*). Briefly, breastmilk samples were diluted 1:1 with sterile phosphate buffered saline (PBS; Life Technologies), centrifuged, washed, and resuspended in PBS prior to centrifugation three times. The resultant breastmilk cell pellet was then processed for single-cell RNA sequencing (scRNAseq), quantitative real-time PCR (RT-qPCR), or flow cytometry.

### Cell viability and flow cytometry analysis

Cell pellets were resuspended in 100 μl of 10% fetal bovine serum (FBS; VWR) and incubated with either Trypan Blue (Invitrogen) or LIVE/DEAD™ Fixable Yellow stain (Invitrogen) for 5 minutes at room temperature and 30 minutes at 37°C, respectively. Fixable Yellow samples were subsequently fixed using Flow Cytometry Fixation Buffer (R&D Systems), centrifuged, and resuspended in 10% FBS in PBS. Trypan Blue cell viability was measured using Countess™ II Automated Cell Counter (ThermoFisher). For flow cytometry, samples were fixed in Fixation Buffer for 30 minutes at 4°C, washed with Permeabilization Buffer (R&D Systems), centrifuged, and incubated with fluorescent antibodies (Table S2) in Permeabilization Buffer at 4°C for 1 hour. Samples were then washed with 10% FBS in PBS, centrifuged, and resuspended in 10% FBS in PBS. Appropriate negative internal controls were used. Data acquisition was done with a NovoCyte® 3000 (ACEA Biosciences) and data analysis with NovoExpress.

### Gene expression analysis by RT-qPCR

Cell pellets were lysed in Buffer RLT, mixed with an equal volume of 100% ethanol, and the RNA was extracted with RNeasy mini kit (Qiagen). cDNA was generated from 1500 ng of RNA using High-Capacity cDNA Reverse Transcription Kit (Applied Biosystems). RT-qPCR was performed using SYBR™ Select Master Mix (Applied Biosystems) on a ViiA 7 Real-Time PCR System (Applied Biosystems). Primer sequences are presented in Table S3.

### Single-cell RNA sequencing

Cells were processed with Chromium Next GEM Single Cell 3’ GEM Library and Gel Bead Kit v 3.1 (10X Genomics) according to manufacturer instructions. Briefly, cells were diluted in master mix containing reverse transcription reagents and primer and were transferred to the ChromiumChip with gel beads and partitioning oil for preparation of nanoliter-scale Gel Bead-In-Emulsions (GEMs). Reverse transcription produced cDNA with a cellular 10X barcode and unique molecular identifier (UMI), and the cDNA was recovered with Dynabeads MyOne SILANE magnetic beads. The cDNA was then subjected to 8-14 cycles of amplification before clean-up using SPRIselect Reagent (Beckman). Total cDNA yield was calculated following quality assessment (High Sensitivity D5000 ScreenTape, Agilent) and fluorometric quantitation (Qubit, ThermoFisher). Libraries were constructed by enzymatic fragmentation, end repair and A-tailing according to the manufacturer’s instructions. Two rounds of clean-up were performed: First, a double sided SPRIselect Reagent (Beckman) prepared samples for adapter ligation. Then, another round of magnetic bead clean-up was performed before sample indexing PCR for 5-16 cycles depending on the cDNA input for library construction. A further double sided size selection produced libraries of 400-600 bp, which were quantified for paired end sequencing (26 bp x 98 bp). The 10X single cell RNA libraries quality was checked using Fragment Analyzer (NGS Fragment Kit, Agilent) and they were quantified by qPCR (Kapa qPCR quantification kit, Kapa biosystem). The libraries were then normalized, pooled, and sequenced on NovaSeq6000 platform (Illumina) with S1 100 cycles kit (read1: 28 bp; read2: 91 bp).

### scRNAseq analysis

All reads were processed using the 10X cellranger pipeline. Briefly, reads were demultiplexed, using the cellranger mkfastq function. Demultiplexed reads from all lanes were aligned to the GRCh38 genome, followed by filtering, barcode counting, and UMI counting using the cellranger count function. After constructing the gene expression matrix, cells were processed using Seurat and Monocle packages. Cells were filtered to remove low quality cells (< 400 genes), doublets (> 5,000 genes), and cells with high mitochondrial fraction (> 25%). Cells were assigned cell cycle scores based on the presence of S, G1, and G2M genes. The filtered subset was normalized and scaled using Seurat’s SCT pipeline. Mitochondrial percentage, cell cycle scores, gene counts, and UMI counts were regressed out. Samples from all three donors were integrated using Seurat’s SCTransform integration pipeline. Highly variable genes from all 26,922 cells were used as features for Principal Component Analysis (PCA) using the RunPCA function. The first 50 principal components were used for UMAP dimensionality reduction using the RunUMAP function. The integrated RNA counts from the Seurat object along with the first 50 principal components and the UMAP coordinates were transferred to a cell_data_set for clustering and trajectory inference using monocle3. Cells were clustered using the Leiden algorithm in monocle using the cluster_cells function. For trajectory inference, the principal graph was learned using the learn_graph function. Cells were ordered in pseudotime by choosing the root node using the order_cells function. A node in the epithelial cell cluster with high expression of KRT7 and KRT18 was chosen as the root node. Gene dynamics as a function of pseudotime were analyzed using the plot_genes_in_pseudotime function. Finally, the clusters from monocle3 were added to the Seurat object’s metadata. Differentially expressed genes were identified using the MAST algorithm with the FindAllMarkers function.

### Statistical analysis

All data are represented as mean ± standard error of the mean (SEM). Statistical analysis was carried out using Prism 8 software (GraphPad) using Student’s t-test and one-way ANOVA with Bonferroni’s post hoc tests. A significant difference was defined as P < 0.05.

## H2: Supplementary Materials

**Fig. S1.**
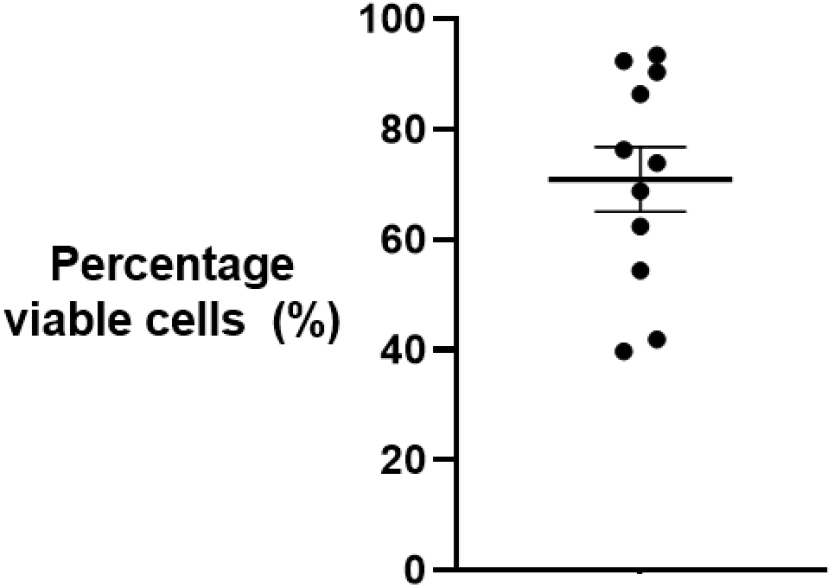
– Cells isolated from the breastmilk of 12 unique donors were 40 to 95% viable with an average viability of 70%. Viability was measured using flow cytometry LIVE/DEAD™ Fixable Yellow stain (79.2% ± 6.9%; N = 10).

**Fig. S2.**
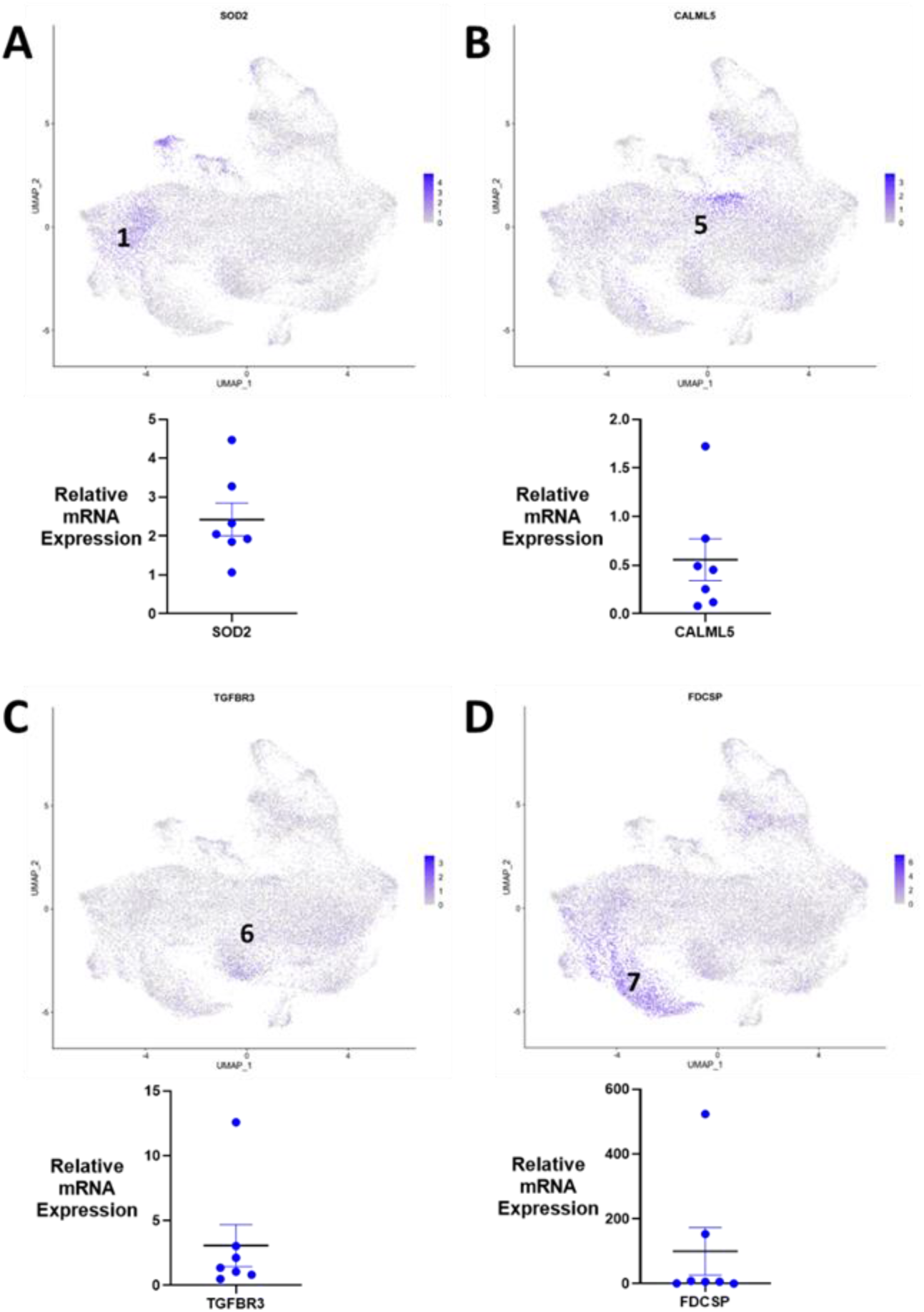
–Cluster marker mRNA expression in a larger donor pool is relatively consistent (A) UMAP localization and corresponding mRNA expression of epithelial cell marker, progenitor cells (*SOD2*; cluster 1/5) and differentiating cell marker (*CALML5*: cluster 5) are consistently expressed across multiple donor samples. Markers for specialized sub-types are enriched in specific donors (C; *PTPRF*+ lactocytes) and (D; chemotactic epithelial cells). UMAP analysis (N = 3) and RT-qPCR (N = 7).

**Table S1.** Donor characteristics and experimental groups

| Donor ID | Weeks Postpartum | Health status | Viability assay | Cell type flow cytometry | Cell type qPCR | Stem cell flow cytometry | Stem cell qPCR | scRNAseq | RNAseq qPCR |
| --- | --- | --- | --- | --- | --- | --- | --- | --- | --- |
| 2.1 | 18 | H |  | Y | Y | Y | Y |  | Y |
| 2.2 | 28 | H |  | Y |  | Y |  |  |  |
| 2.3 | 31 | H | Y |  |  |  |  | Y |  |
| 4 | 38 | H |  | Y |  | Y |  |  |  |
| 5 | 83 | H | Y | Y |  | Y |  |  |  |
| 6.1 | 12 | H |  | Y | Y | Y | Y |  | Y |
| 6.2 | 19 | H |  | Y |  | Y |  |  |  |
| 6.3 | 20 | H | Y |  |  |  |  | Y |  |
| 8.1 | 39 | H | Y | Y |  | Y |  |  |  |
| 8.2 | 44 | H |  | Y |  | Y |  |  | Y |
| 9 | 21 | M | Y | Y |  | Y |  |  |  |
| 11 | 11 | M | Y | Y | Y | Y | Y |  | Y |
| 12.1 | 29 | H |  | Y |  | Y |  |  |  |
| 12.2 | 32 | H |  | Y |  | Y |  |  |  |
| 12.3 | 33 | H | Y |  |  |  |  | Y |  |
| 13 | 92 | H | Y | Y | Y | Y | Y |  | Y |
| 17 | 6 | S | Y |  |  |  |  |  |  |
| 18 | 5 | H |  | Y | Y | Y | Y |  | Y |
| 19 | 36 | M |  |  |  |  |  |  |  |
| 21 | 40 | H | Y | Y | Y | Y | Y |  | Y |

Health status: H = healthy mother and infant, M = recent mastitis infection, and S = infant currently sick.

**Table S2.** Flow cytometry antibodies used in analysis

| Marker | Fluorophore | Company | Cat. No |
| --- | --- | --- | --- |
| EpCAM | PE-Cy7 | eBioscience | 25-9326-42 |
| CD45 | Alexa Fluor® 488 | R&D Systems | FAB1430G |
| Vimentin | APC | Invitrogen | MA528601 |
| CK18 | FITC | Invitrogen | MA110327 |
| aSMA | Alexa Fluor® 594 | R&D Systems | IC1420T-100 |
| SOX2 | eFluor® 660 | eBioscience | 50-9811-82 |
| TRA-1-60 | DyLight® 488 | Invitrogen | MA1023D488X |
| Nanog | PE | Invitrogen | PA546891 |
| SSEA4 | PerCP-eFluor® 710 | eBioscience | 46-8843-42 |
| CD49f | eFluor® 450 | eBioscience | 48-0495-82 |

**Table S3.**
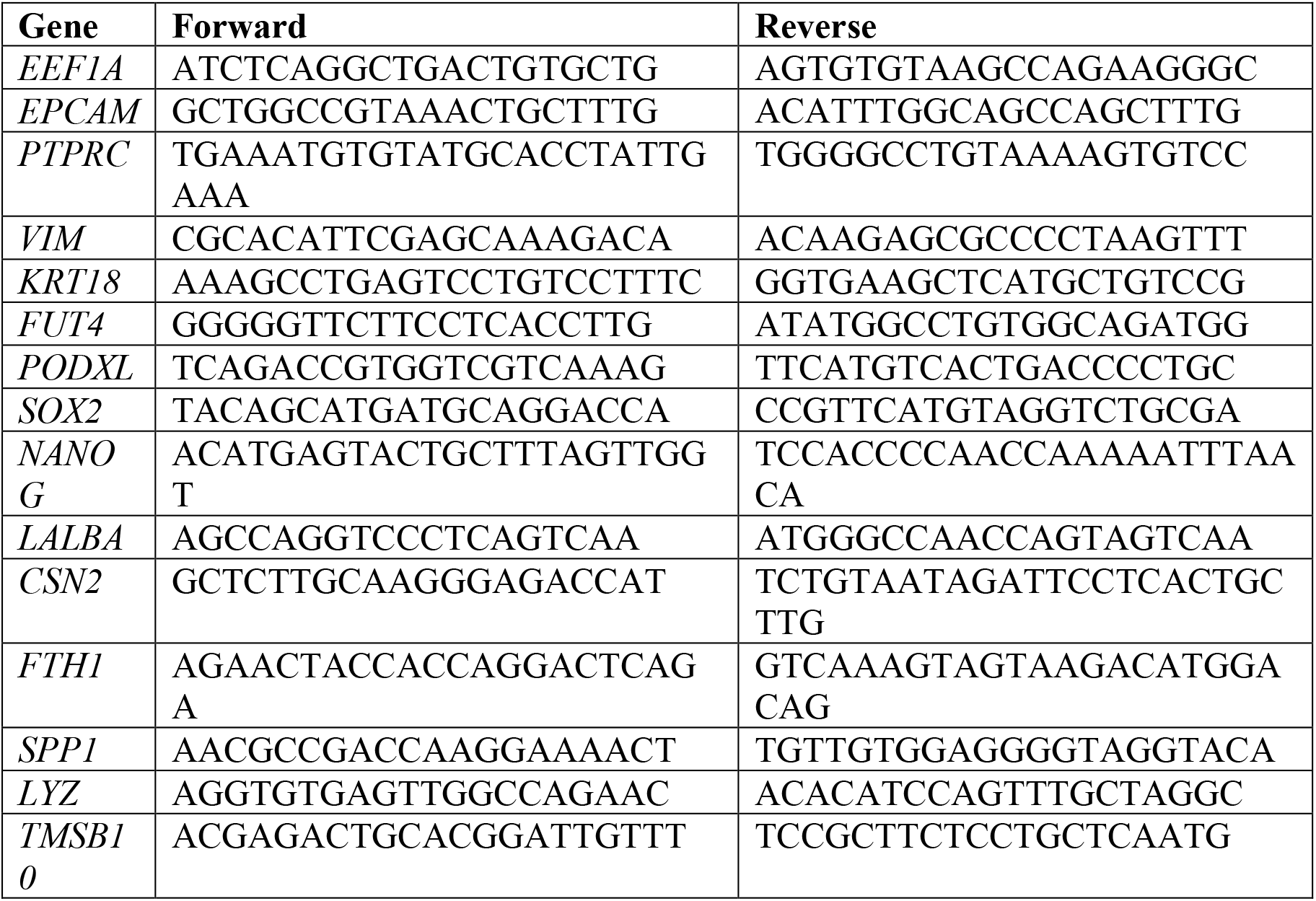
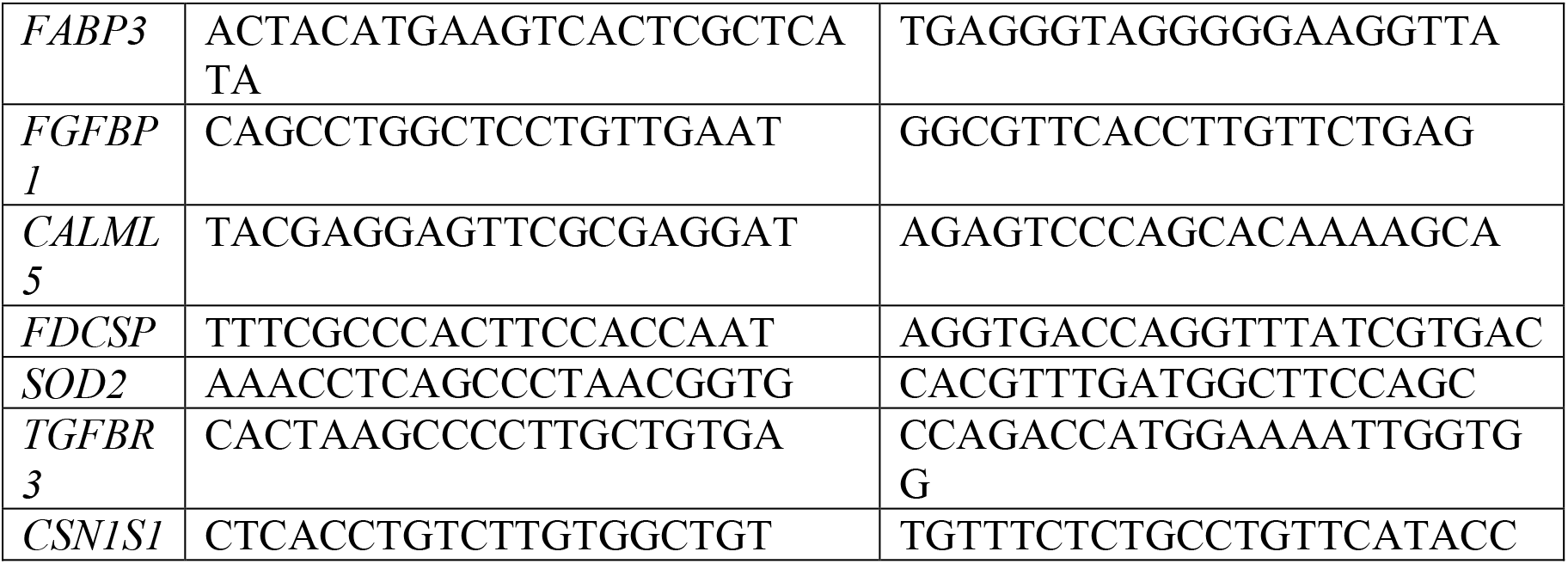
Primer sequences for RT-qPCR

## Acknowledgments

This project used the University of Pittsburgh HSCRF Genomics Research Core for RNAseq prep and library prep, and we thank H. Monroe and D. Hollingshead for their assistance. This work was supported by the UPMC Genome Center with funding from UPMC’s Immunotherapy and Transplant Center for single cell sequencing, and we thank Y. Pan for their assistance.

## General

We thank our donors for providing breastmilk samples and making this work possible.

## Author contributions

J.P.G and K.A.W conceived and designed the experiments. R.D. and T.H.S. recruited donors. J.P.G., N.C. and K.C.F. performed the experiments. J.P.G., N.C. and K.A.W. analyzed the data. J.P.G., N.C., R.D., K.C.F., T.H.S., and K.A.W. contributed to writing and editing the manuscript.

## Data and materials availability

All data needed to evaluate the conclusions in the paper are present in the paper. Additional data related to this paper may be requested from the authors.

